# Chemical-genetic interaction mapping links carbon metabolism and cell wall structure to tuberculosis drug efficacy

**DOI:** 10.1101/2021.04.08.439092

**Authors:** Eun-Ik Koh, Peter O. Oluoch, Nadine Ruecker, Megan K. Proulx, Vijay Soni, Kenan C. Murphy, Kadamba G. Papavinasasundaram, Charlotte J. Reames, Carolina Trujillo, Anisha Zaveri, Matthew D. Zimmerman, Roshanak Aslebagh, Richard E. Baker, Scott A. Shaffer, Kristine M. Guinn, Michael Fitzgerald, Véronique A. Dartois, Sabine Ehrt, Deborah T. Hung, Thomas R. Ioerger, Eric Rubin, Kyu Y. Rhee, Dirk Schnappinger, Christopher M. Sassetti

## Abstract

Current chemotherapy against *Mycobacterium tuberculosis* (*Mtb*), an important human pathogen, requires a multidrug regimen lasting several months. While efforts have been made to optimize therapy by exploiting drug-drug synergies, testing new drug combinations in relevant host environments remains arduous. In particular, host environments profoundly affect the bacterial metabolic state and drug efficacy, limiting the accuracy of predictions based on *in vitro* assays alone. In this study, we utilize conditional *Mtb* knockdown mutants of essential genes as an experimentally-tractable surrogate for drug treatment, and probe the relationship between *Mtb* carbon metabolism and chemical-genetic interactions (CGI). We examined the anti-tubercular drugs isoniazid, rifampicin and moxifloxacin, and found that CGI are differentially responsive to the metabolic state, defining both environment-independent and –dependent interactions. Specifically, growth on the *in vivo*-relevant carbon source, cholesterol, reduced rifampicin efficacy by altering mycobacterial cell surface lipid composition. We report that a variety of perturbations in cell wall synthesis pathways restore rifampicin efficacy during growth on cholesterol, and that both environment-independent and cholesterol-dependent *in vitro* CGI could be leveraged to enhance bacterial clearance in the mouse infection model. Our findings present an atlas of novel chemical-genetic-environmental interactions that can be used to optimize drug-drug interactions as well as provide a framework for understanding *in vitro* correlates of *in vivo* efficacy.

**Significance:** Efforts to improve tuberculosis therapy include optimizing multi-drug regimens to take advantage of drug-drug synergies. However, the complex host environment has a profound effect on bacterial metabolic state and drug activity, making predictions of optimal drug combinations difficult. In this study, we leverage a newly developed library of conditional knockdown *Mycobacterium tuberculosis* mutants in which genetic depletion of essential genes mimics the effect of drug therapy. This tractable system allowed us to assess the effect of growth condition on predicted drug-drug interactions. We found that these interactions can be differentially sensitive to the metabolic state and select *in vitro*-defined interactions can be leveraged to accelerate bacterial killing during infection. These findings suggest new strategies for optimizing tuberculosis therapy.

## Introduction

The current chemotherapeutic regimen for tuberculosis (TB) is the product of many decades of basic and clinical research. Since the first trials of streptomycin monotherapy in 1948 were rapidly followed by the emergence of antibiotic resistant clones (1), multidrug regimens to both suppress resistance and accelerate bacterial killing have become standard. The current regimen used against drug-sensitive strains of *Mycobacterium tuberculosis* (*Mtb*) consists of four antibiotics, isoniazid (INH), rifampicin (RIF), pyrazinamide (PZA), and ethambutol (EMB), and was optimized in a series of clinical trials in the 1970s (2, 3). While this “short-course regimen” has been credited with curing over 50 million patients, its delivery is complicated by the need for 6-9 months of drug administration (4). Furthermore, even in clinical trial settings where the delivery of this extended regimen is assured, 5-10% of patients fail therapy (5). The frequent transmission of antibiotic resistant *Mtb* strains has further complicated TB treatment options, requiring the use of less optimized drug combinations that are administered for even longer periods (6).

The factors that necessitate prolonged therapy are complex and specific to the infection environment. While antibiotics such as INH and RIF cause rapid cell death *in vitro*, their antimicrobial activities are much slower during infection (7). Limited drug penetration into *Mtb*-containing tissue lesions may reduce the efficacy of some drugs. For example, the intralesional RIF concentrations are significantly lower than what is achieved in the plasma (8, 9), and clinical studies suggest that increased RIF dosing improves bacterial clearance (10, 11). However, drug penetration is unlikely to fully account for the reduced antibiotic efficacy in host tissue, as this complex and stressful environment has also been shown to alter the physiology of the pathogen to induce a drug-tolerant state (12–14). While a number of host immune-related stresses may be involved in this process, simple changes in macronutrient availability can have important consequences. *Mtb* has access to a mixture of glycolytic carbon sources, fatty acids and cholesterol in host tissue (15–17), and altering the availability of these carbon sources *in vitro* can change the efficacy of anti-tubercular compounds (18–23). A role for differential carbon catabolism in determining drug efficacy is also supported by the identification of natural genetic variants in clinical *Mtb* isolates that enhance drug tolerance by altering either glycerol, lipid, or sterol catabolism (24, 25). The profound effect of the host environment on bacterial metabolic state and drug activity makes it difficult to predict the ultimate efficacy of an antibiotic regimen based on *in vitro* assays alone.

The most advanced efforts to accelerate TB therapy involve optimizing multi-drug regimens to take advantage of drug-drug synergies. These pharmacological interactions can improve therapy by increasing drug exposure, decreasing MIC, or enhancing the maximal effect of the treatment (26). However, despite the demonstrated benefits of synergistic regimens in many therapeutic realms, the combinatorial burden of testing all potential multi-drug combinations remains cumbersome. While recent advances to more efficiently predict synergies *in vitro* have proven valuable (27, 28), it remains unclear whether the environmental influences that alter the efficacy of individual drugs will also influence drug-drug interactions (29, 30).

In this study, we leveraged a newly developed genetic resource to understand the effect of growth conditions on potential drug-drug interactions. This “hypomorph library” consists of individually DNA barcoded strains in which individual essential genes are tagged with the DAS+4 sequence that targets proteins for degradation upon inducible induction of the SspB adapter (31). Graded *sspB* expression produces different degrees of depletion to model the effect of partial chemical inhibition (32). This approach allowed for highly parallel assessment of changes in TB drug sensitivity of individual hypomorphic mutants under different growth conditions; creating a drug-specific chemical-genetic interaction (CGI) profile (33). We report that drugs of distinct classes produce interactions that are differentially sensitive to the environment, and that both condition-independent and –dependent CGIs can be identified for all drugs tested. In particular, RIF efficacy was impaired as a result of cell surface lipid alterations that occur during growth in cholesterol, and the cell could be resensitized through a condition-specific interaction with the cell wall synthetic machinery both *in vitro* and in the mouse lung. These observations provide a compendium of chemical-genetic interactions that can be exploited to enhance therapy and begin to define *in vitro* correlates of *in vivo* efficacy.

## Results

### Genetic strategy to define essential bacterial functions that alter drug efficacy *in vitro*

Host environmental factors alter antibiotic efficacy against TB (12, 24, 34, 35). To determine if carbon source availability specifically affects antibiotic efficacy, we used the *Mtb* strain H37Rv in minimal medium with glycerol, acetate or cholesterol as sole carbon sources in a growth inhibition assay with the TB drugs isoniazid (INH), rifampicin (RIF) and moxifloxacin (MOX). As these carbon sources support different growth rates, endpoint-based antibiotic activity measurements such as MIC assays can be difficult to interpret. Therefore we quantified growth rates in each carbon source over multiple concentrations of antibiotics to determine the concentration that decreased the growth rate by 50% (GR_50_) (24, 36). Growth of individual strains was monitored during exponential phase (OD_600_ 0.20-0.25) and the growth rate was determined by fitting an exponential curve **(Fig1A)**. We found different carbon sources to alter the GR_50_ of these drugs in distinct ways. For INH, glycerol-dependent growth increased efficacy, relative to the non-glycolytic carbon sources. In contrast, growth in cholesterol was found to decrease RIF efficacy, relative to the other conditions. Media composition had a more modest effect on MOX, altering GR_50_ by only approximately 2-fold **(Fig1B)**. These findings show that carbon metabolism plays an important role in antibiotic efficacy and has distinct effects on different drugs.

**Figure 1.**
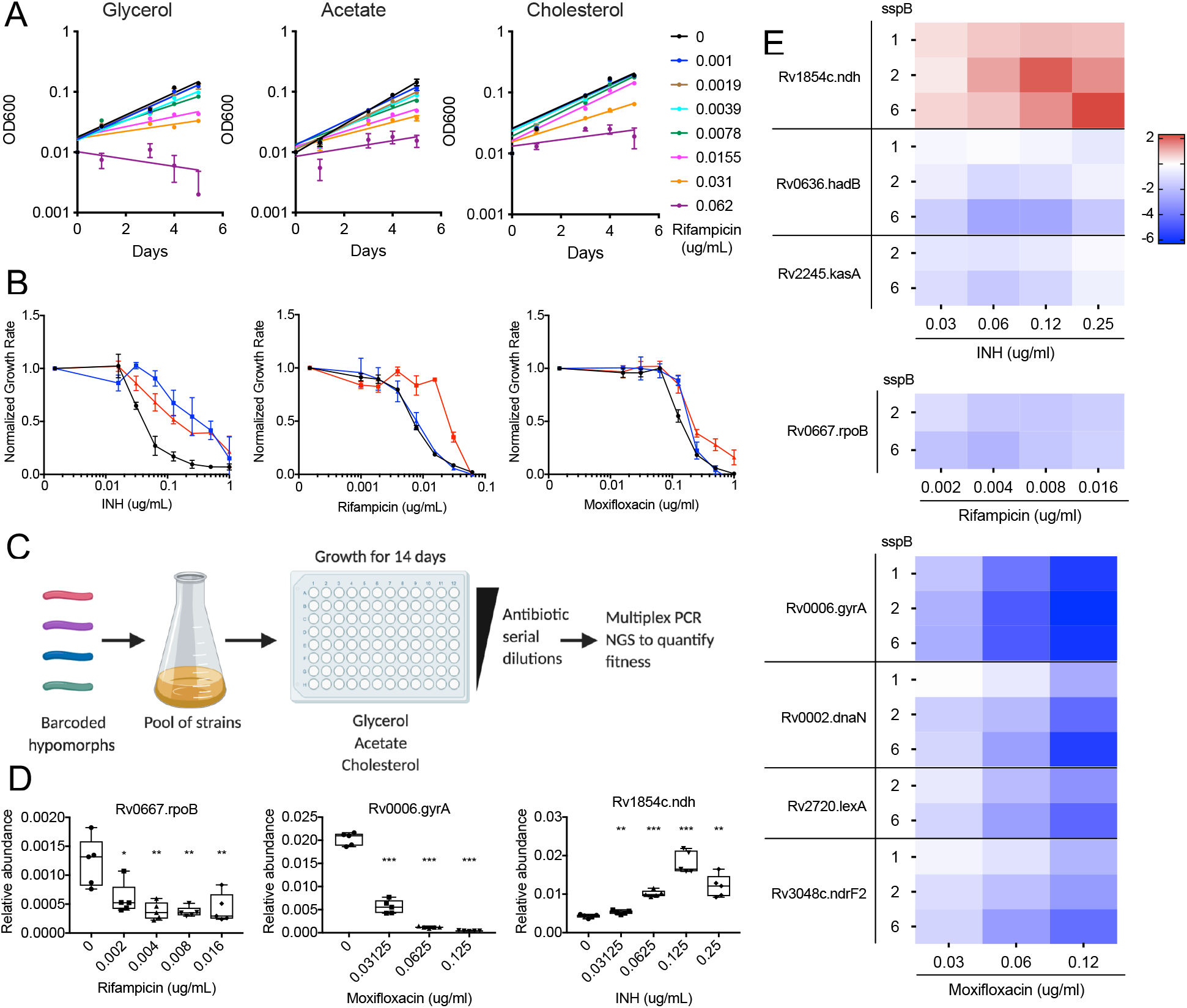
Genetic strategy to define essential bacterial functions that alter drug efficacy *in vitro*. **(A)** Growth of H37Rv in minimal media with glycerol, acetate and cholesterol as the sole carbon source and increasing concentrations of RIF. Results shown as mean from 3 biological replicates with standard deviations. **(B)** Normalized growth inhibition of WT *Mtb* across increasing concentrations of INH, RIF and MOX in minimal media with glycerol (black), acetate (blue) and cholesterol (red) as sole carbon sources. Results shown as means from 3 biological replicates with standard deviations. **(C)** Barcoded hypomorph mutants were pooled and grown in 96-well plates containing minimal media with glycerol, acetate or cholesterol as the sole carbon source for 14 days. Antibiotics were added to individual wells as well as untreated controls. Chromosomal barcodes were PCR amplified and pooled for Illumina Next Generation Sequencing (NGS). Barcodes were analyzed to quantify changes in fitness of individual strains from different conditions. **(D)** Boxplot representing changes in relative abundances of *rpoB*, *gyrA* and *ndh* mutants grown in glycerol during RIF, MOX and INH treatment respectively. The highest drug concentration tested for RIF, MOX and INH represent the concentration at which most of the strains in the library were inhibited. Data represents 5 biological replicates. Significance was calculated using unpaired t-test and compared to untreated conditions, \**p*<0.05, \*\**p*<0.01, \*\*\**p*<0.001. **(E)** Heat map representing changes in fitness of individual mutants, shown as Log2 fold change, during INH, RIF and MOX treatment in glycerol growth conditions. Mutants were chosen from previous association with respective antibiotic. sspB numbers denote *sspB* expression level of mutant. Results shown as means from 5 biological replicates.

To understand the mechanisms linking drug efficacy and metabolism, we utilized a barcoded-hypomorph library consisting of 465 essential genes to identify CGI with these drugs (32). The library was grown in minimal media with glycerol, acetate or cholesterol, and antibiotics were added at concentrations ranging from 0.05X to 1X GR_50_ for each. Bacteria were subjected to these conditions in 96-well plates for 14 days, at which point the relative abundance of individual mutants was assessed through multiplex PCR and Next-Generation Sequencing **(Fig1C, S.Table 1)**. Relative abundances of individual mutants were calculated by normalizing the mutant’s barcode count to the total barcode count and comparing the relative abundance of each mutant in different drug concentrations with the untreated control samples.

We initially investigated the validity of this dataset by determining if genetic inhibition of each drug’s target produced the expected CGI with that compound in standard glycerol-dependent growth conditions. RIF and MOX inhibit the RNA polymerase subunit, RpoB or the DNA gyrase subunit, GyrA, respectively. As expected, *rpoB* and *gyrA* hypomorphic mutants were hypersensitive to RIF or MOX, and the CGI increased with drug concentration **(Fig1D)**. The major target of INH, InhA, was not present in our mutant pool. However, the library did contain a *ndh* mutant. Deficiency in this NADH-dehydrogenase is known to decrease INH efficacy, likely due to the inefficient formation of the active INH-NAD adduct (37). Consistent with these observations, we found that the *ndh* hypomorphic mutant showed an INH dose-dependent increase in abundance in our dataset **(Fig1D)**.

The hypomorph library was constructed to contain up to five different versions of each DAS+4 tagged mutant, which express different levels of *sspB* and therefore produce graded levels of protein depletion (31, 38). To investigate the relationship between target protein abundance and phenotype, we correlated the strength of *sspB* expression with drug sensitivity for a number of genes that were expected to alter antibiotic efficacy **(Fig1E)**. For RIF, we concentrated on *rpoB*. For INH, we examined *ndh* as well as genes associated with mycolate biosynthesis, and found *kasA* and *hadB* mutants to be significantly hypersensitive to increasing INH concentrations. For MOX, in addition to *gyrA*, depletion of proteins associated with DNA integrity or deoxynucleotide production, *dnaN, lexA* and *ndrF2*, showed significantly decreased fitness over increasing MOX concentrations. For some mutants, particularly those involved in DNA metabolism, we noted a dose-dependent effect of *sspB* expression, with larger phenotypes corresponding to greater degrees of depletion. However, this was not a universal phenomenon, and we often found that independent mutants expressing different *sspB* levels produced consistent phenotypic effects. The observed differences in sensitivity to protein depletion likely reflect the unique biochemistry of each pathway (39). As a result, we considered each mutant corresponding to a target protein independently in the following analyses.

### CGI profiles are determined by the combination of carbon source and drug

To understand the relative importance of carbon metabolism and drug on bacterial physiology, we compared the “fitness profiles” of each condition, which represented the relative fitness of each mutant strain. We first selected the fold change in abundance (log2FC) values of every mutant at 0.65X GR_50_ drug concentration that showed a statistically significant fitness difference when at least one treated sample was compared to the untreated control (918 mutants, Q <0.05). 0.65X GR_50_ drug concentration displayed the largest number of mutants with a statistically significant fitness difference across most conditions. Upon hierarchical clustering of both genes and conditions, we found three major condition clusters that corresponded to each antibiotic **(Fig2A)**. These condition clusters were differentiated by two distinct clusters of mutants; one large gene set differentiated MOX from the other drugs (Fig 2A bottom cluster), and differential fitness effects on a second smaller gene set differentiated INH and RIF (Fig2A top cluster). Within each condition cluster, fitness effects due to carbon source were apparent.

**Figure 2.**
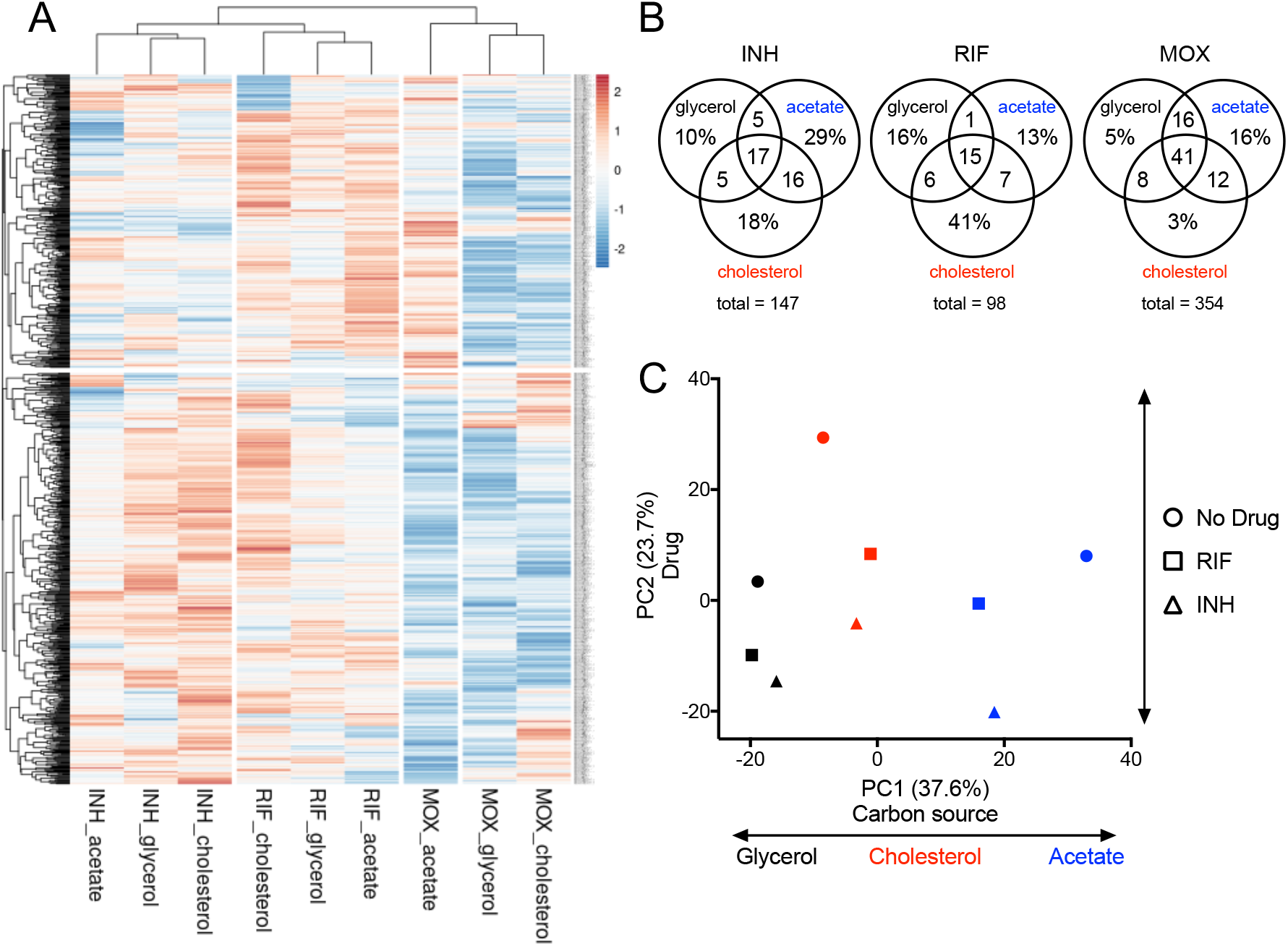
Carbon sources mediate antibiotic-genetic interactions of essential bacterial functions. **(A)** Heat map of hypomorph mutants shown as Log2 fold change of 0.65X GR_50_ of INH, RIF and MOX against untreated controls. Results shown as means from 5 biological replicates. Each antibiotic (INH, RIF and MOX) in single carbon source (glycerol, acetate and cholesterol) conditions were compared using hierarchical clustering methods. **(B)** Venn diagrams showing the total number of genes with log2FC > −1 and *p*<0.05 at 0.65X GR_50_ of INH, RIF and MOX, and the % of genes associated with specific conditions. (**C)** Relative abundance datasets of the hypomorph libraries from glycerol (black), acetate (blue) and cholesterol (red) growth conditions with 0.65X GR_50_ INH (triangle) and RIF (square) treatment as well as untreated controls (circle) were examined using principal component analysis. Units shown on axes are arbitrary values in principal component space.

To catalog the genes that differentiate these conditions, we compared each drug-treated sample (0.65X GR_50_) with the corresponding untreated control and chose genes with log2FC < −1 and *p* < 0.05 for each carbon source. For all antibiotics tested, there were mutants that showed decreased fitness across all carbon sources as well as those associated with a single carbon source **(Fig2B)**. Notably, MOX produced the largest number of mutants with decreased fitness in a condition-independent manner, while INH and RIF produced the largest number of mutants with decreased fitness during acetate and cholesterol-dependent growth respectively. To more quantitatively assess the relative contribution of carbon source and drug, we used Principal Component Analysis (PCA) to examine the amount of variance contributed by each, concentrating only on the INH and RIF conditions, which produced the most condition-specific effects. PC1 aligned to distinguish the carbon sources, while PC2 differentiated among drug treatments **(Fig2C)**. The first and second principal components accounted for 37.6% and 23.7% respectively, of the overall variance across the 9 data sets, indicating that drug and carbon source played a relatively equal role in shaping bacterial physiology.

### Condition-independent CGI can be reproduced with individual mutants

Genes found to alter antibiotic efficacy from the high-throughput hypomorph analysis were validated using individual mutant strains. For initial validation studies, we concentrated on mutants predicted to show drug-hypersensitivity under all media conditions. We also replaced acetate with butyrate, as butyrate provided relatively enhanced growth that allowed for robust validation. As these hypomorph mutants target essential genes that may alter growth, we compared the relative growth rates of individual mutants to WT in different carbon sources **(SFig1)**. Variability in growth rates of mutants in different growth conditions led to the utilization of the growth rate inhibition method described above to determine the RIF MIC. The repression of a drug’s target is thought to increase sensitivity to inhibition by reducing the fraction of the target that needs to be inhibited to reduce growth (40). As such, we anticipated that drug target inhibition would result in condition-independent effects. This prediction was verified using a *rpoB* hypomorphic mutant, which our screening data predicted to increase RIF efficacy in all carbon sources **(Fig3A)**. As such, we verified that this degree of RpoB depletion decreased the RIF MIC by 4 to 16-fold in each of the three media conditions **(Fig3A)**. In addition to known drug targets, a number of additional genes were predicted to enhance RIF efficacy in a carbon source-independent manner, including the transcriptional regulator, Rv0472c. This gene has been named, mycolic acid desaturase regulator (*madR*), based on its specific effect on the mycolate-modifying *desA1* and *desA2* genes (41). As we found for RpoB, MadR depletion sensitized the bacterium to RIF in all tested carbon sources in the single strain assay **(Fig3B)**. Finally, to ensure the specificity of these CGI, we investigated whether they could be reversed by specific metabolite complementation. As predicted by the screen, depletion of the lysine biosynthetic enzyme, LysA, produced RIF hypersensitivity in both glycerol and cholesterol media, and we found this phenotype was largely reversed by supplementation with 3mM lysine **(Fig3C)**. These data support the predictions of the primary screen, and demonstrate that certain CGI are relatively unaffected by carbon source.

**Figure 3.**
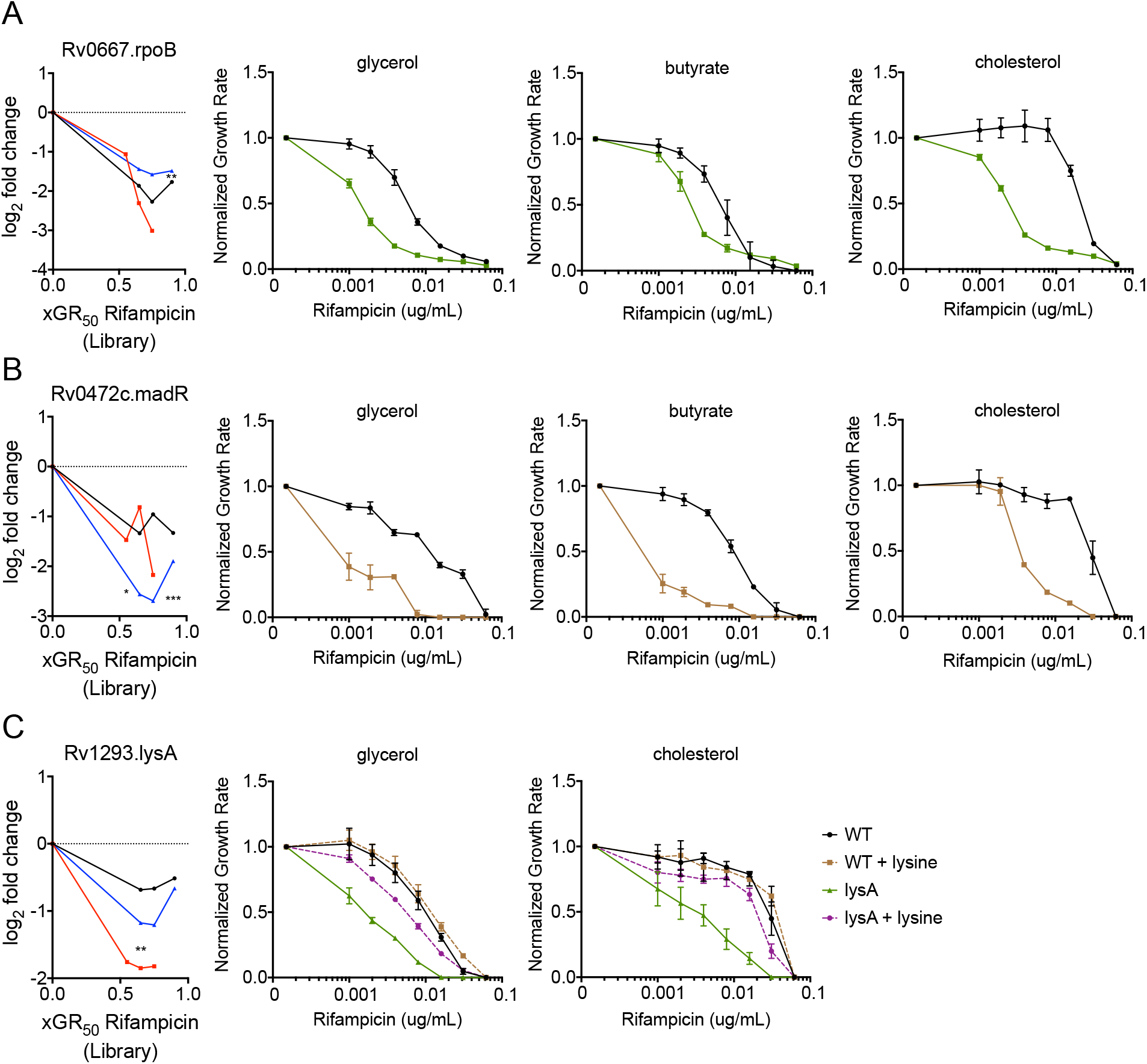
Condition-independent chemical-genetic interactions can be reproduced with individual mutants. **(A)** RpoB depletion shows decreased fitness compared to the library over increasing concentrations of RIF for glycerol (black), acetate (blue) and cholesterol (red). Normalized growth inhibition assay shows *rpoB* hypomorph mutant (green) has decreased RIF MIC compared to WT (black) across multiple carbon conditions. **(B)** MadR depletion shows decreased fitness compared to the library over increasing concentrations of rifampicin for glycerol (black), acetate (blue) and cholesterol (red). Normalized growth inhibition assay shows *madR* hypomorph mutant (brown) has decreased RIF MIC compared to WT (black) across multiple carbon conditions. **(C)** LysA depletion shows decreased fitness compared to the library over increasing concentrations of rifampicin for glycerol (black), acetate (blue) and cholesterol (red). Normalized growth inhibition assay shows *lysA* hypomorph mutant (green) has decreased RIF MIC compared to WT (black), which can be reversed by 3mM lysine supplementation (purple). Library results are shown as means from 5 biological replicates, and significance was calculated using unpaired t-test with Benjamini-Hochberg multiple testing correction, \**q*<0.05, \*\**q*<0.01, \*\*\**q*<0.001. Normalized growth inhibition results are shown as means from 3 biological replicates with standard deviations.

### Depletion of cell-wall biosynthetic genes produces condition-specific RIF interactions

As cholesterol represents a primary carbon source for *Mtb* during infection (16, 42), and RIF efficacy is thought to be limited by drug exposure (43–45), we probed the hypomorph fitness profiles for clues to the mechanism underlying the cholesterol-dependent increase in RIF MIC. We found that inhibition of genes associated with different components of the mycobacterial cell wall, including arabinogalactan (*embC* and *aftB*), and mycolic acid (*kasA*) (46, 47) preferentially sensitized the bacterium to RIF during cholesterol growth **(Fig4A)**. These mutants were significantly underrepresented by up to 8-fold upon RIF treatment predominantly in cholesterol growth conditions across multiple *sspB* doses and drug concentrations. To further explore the effect of cell wall inhibition, we tested individual mutants with defects in peptidoglycan (*dapE*), arabinogalactan (*aftB*), and mycolic acid (*hadB*) synthesis (46, 48–50). We first compared the relative growth rates of individual mutants to WT in different carbon sources and found no significant differences for *dapE* and *hadB* hypomorph mutants across all conditions, while *aftB* hypomorph mutant showed modest growth rate increase in glycerol **(SFig2)**. We then found that perturbation in each of these cell wall layers produced only relatively modest effects in glycerol or butyrate media, but all consistently sensitized the bacterium to RIF in cholesterol growth conditions **(Fig4BCD)**. These observations suggest that alterations in cell wall structure might underlie the relationship between cholesterol catabolism and reduced RIF efficacy.

**Figure 4.**
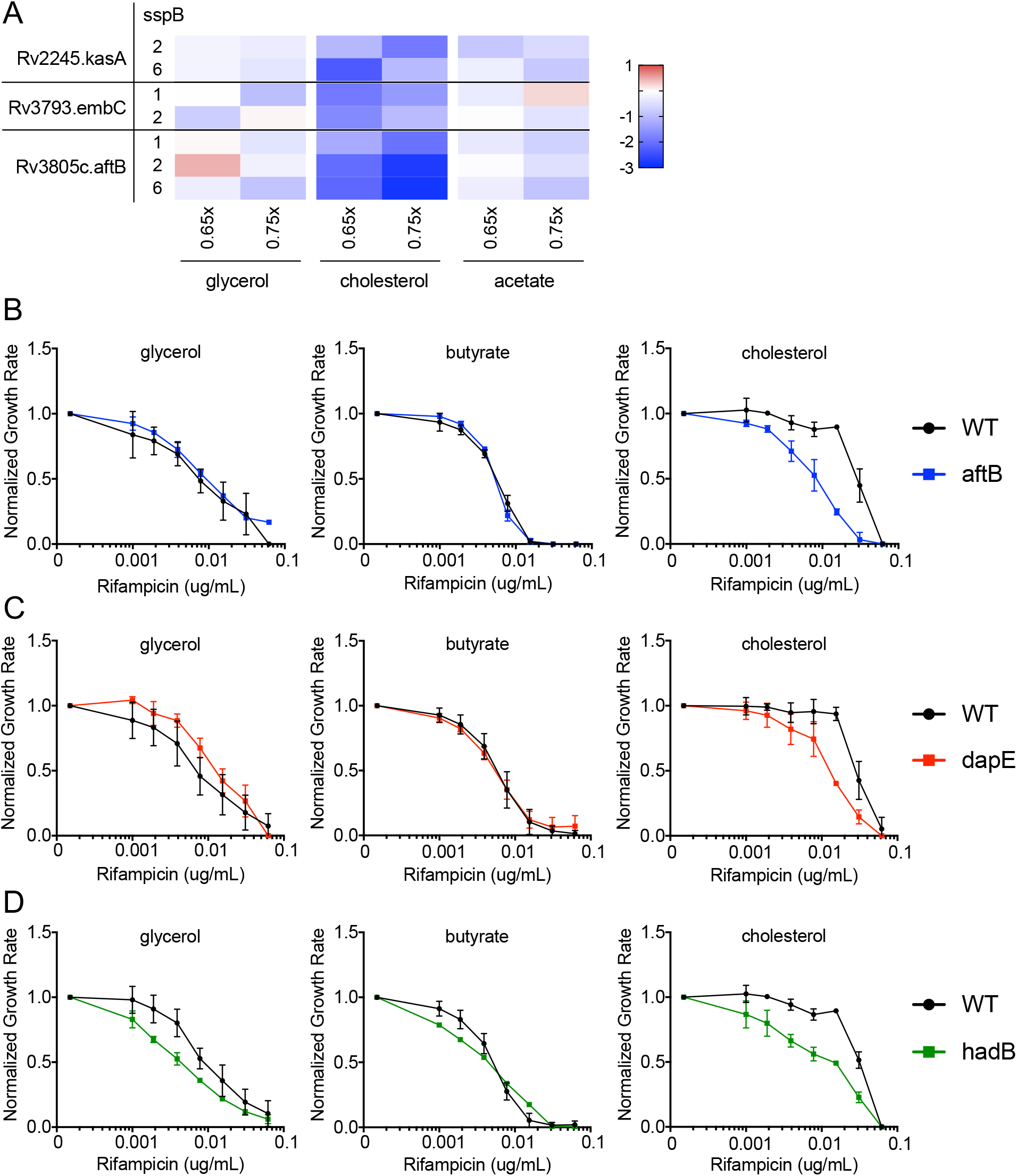
Cell-wall biosynthesis hypomorph mutants show altered RIF efficacy during cholesterol growth conditions. **(A)** Heat map showing hypomorph mutants of select mycobacterial cell-wall biosynthesis genes and pathways with significant changes in fitness during cholesterol growth, shown as Log2 fold change at 0.65X and 0.75X GR_50_ of RIF against untreated controls. Results shown as means from 5 biological replicates. Classification of mutants was done using Mycobrowser. **(B-D)** Normalized growth inhibition of WT and select mutants across increasing concentrations of RIF in minimal media with sole carbon sources. Depletion of AftB (blue), DapE (red) and HadB (green) show decreased RIF MIC compared to WT (black) only during cholesterol growth conditions. Results shown as means from 3 biological replicates with standard deviations.

### RIF efficacy correlates with propionate catabolism and branched chain lipid abundance

*Mtb* catabolizes cholesterol to pyruvate, propionyl-CoA and acetyl-CoA (17, 51). Propionyl-CoA is a precursor of branched-chain lipid synthesis, and catabolism of either cholesterol or propionate increases cellular propionyl-CoA levels causing an increase in both the abundance and chain-length of cell wall lipids, such as diacyltrehalose (DAT), polyacyltrehalose (PAT), and sulfolipid-1 (SL-1) (51–54). To understand whether these alterations in cell wall lipids could be responsible for altering RIF efficacy, we compared RIF GR_50_ values in cells grown in cholesterol to those grown in butyrate media supplemented with increasing concentrations of propionate. Growth rate in cholesterol showed no significant difference compared to both butyrate and butyrate supplemented with propionate **(SFig3)**. We observed that GR_50_ values increased up to 8-fold with increasing propionate levels, and the addition of 0.1X propionate, reflecting a 1:10 ratio of propionate to butyrate (weight:weight), mimicked the effect of cholesterol **(Fig5A)**. To more quantitatively associate branched-chain lipid abundance with drug efficacy, we performed relative quantification of the abundance of SL-1 during growth in butyrate, butyrate and propionate, and cholesterol using mass spectrometry. As expected, we found that growth in either propionate or cholesterol increased both the *m/z* range of SL-1 and its total abundance, and the effect of propionate was dose-dependent in the tested range **(Fig5B)**. This dose-dependency allowed us to correlate changes in sulfolipid abundance with changes in rifampicin efficacy. At multiple propionate concentrations, we calculated the GR_50_ for RIF and the total abundance of SL-1. We found a correlation between these values, as 0.003X propionate supplementation had no effect on either SL-1 abundance or RIF GR_50_, and 0.1X increased both values **(Fig5C)**. These findings implied that the propionyl-CoA derived from cholesterol alters rifampicin efficacy through modification of mycobacterial surface lipids.

**Figure 5.**
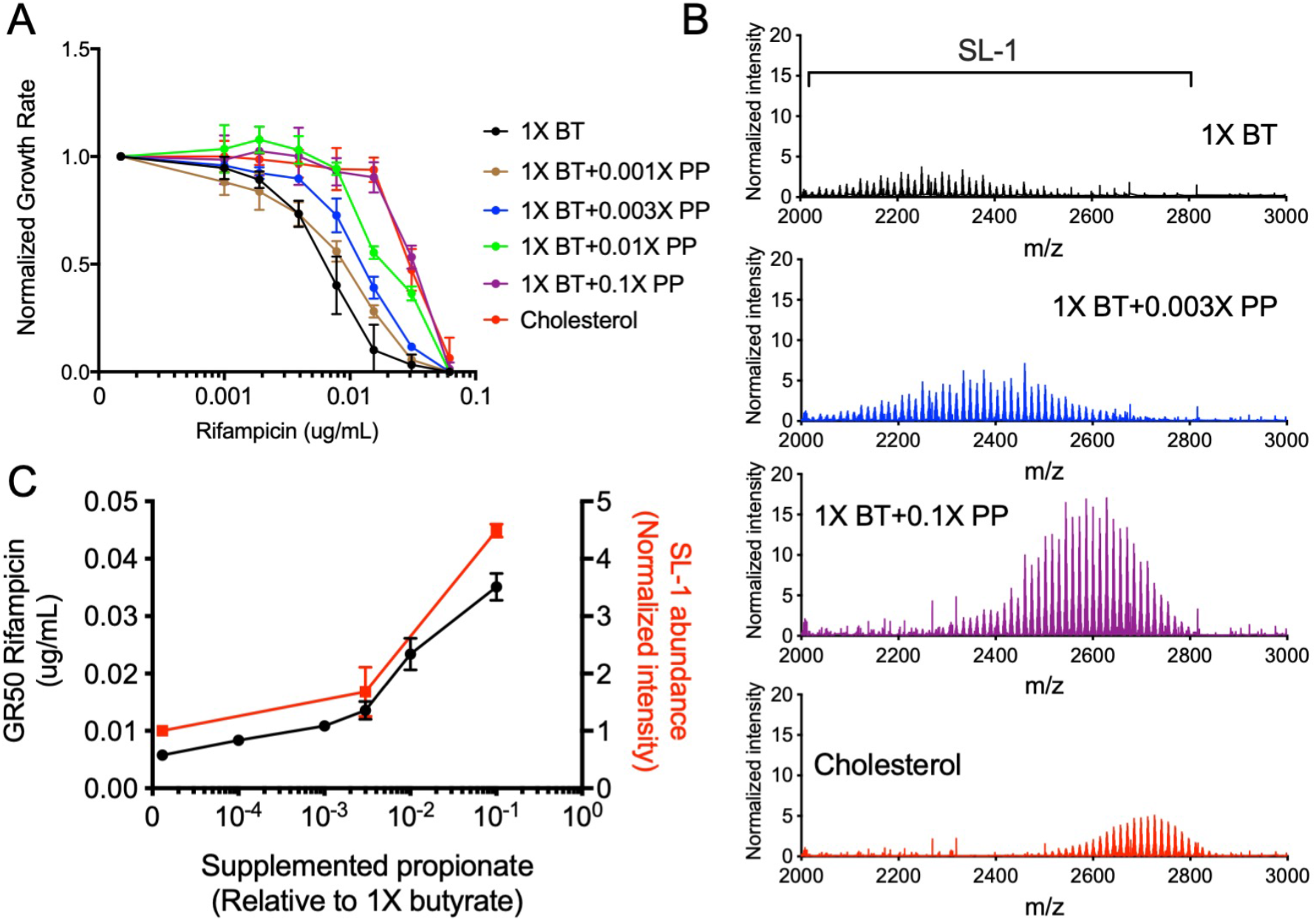
Propionate and cholesterol catabolism decreases RIF efficacy in correlation with increased branched chain lipid length and abundance. **(A)** Normalized growth inhibition of WT across increasing concentrations of RIF in minimal media with 1X butyrate (BT) and corresponding propionate (PP) supplementations as well as cholesterol. Results shown as means from 3 biological replicates with standard deviations. **(B)** Mass spectra of total lipid extract from WT grown in minimal media with 1X butyrate (BT) and corresponding propionate (PP) supplementations as well as cholesterol. MS spectra intensities were normalized using GM2 ganglioside internal standard and cell density. Results shown as representative from 2 independent experiments. Mass range characteristic of sulfolipid-1 is shown. **(C)** Plotted GR_50_ values (black) and relative sulfolipid abundances (red) across increasing propionate levels. Relative sulfolipid abundances were determined by the combined normalized mass spectra intensities from *m/z* 2000-2800 for each growth condition. Results shown as means with standard deviations.

The different branched chain lipid species of *Mtb* are produced by distinct biosynthetic pathways, and inhibiting the synthesis of one lipid can produce a compensatory increase in others (53). Therefore, to assess whether increased lipid abundance was causally related to RIF efficacy, we employed an *Mtb* mutant lacking the PhoPR regulatory system, which is required for the synthesis of multiple branched chain lipid species (55–57). We first compared the relative growth rates of Δ*phoPR* deletion mutant to WT in different carbon sources and found that while Δ*phoPR* deletion mutant showed modest growth rate decrease in butyrate, no significant differences were seen in butyrate with supplemented with propionate and cholesterol conditions **(SFig3)**. In contrast to wild type *Mtb*, a Δ*phoPR* deletion mutant showed little change in rifampicin efficacy with propionate supplementation or cholesterol compared to butyrate growth conditions, and possessed significantly decreased SL-1 levels compared to WT **(Fig6AB)**. Similar to the effects seen with the Δ*phoPR* mutant, cell wall defective hypomorphs (*aftB*, *hadB*, *dapE*) also maintained a relatively consistent RIF GR_50_ in all media conditions, compared to wild type **(Fig6CD)**. These findings suggest that cholesterol catabolism reduces RIF efficacy via a propionyl-CoA driven increase in synthesis of cell surface lipids, and that this effect can be reversed by perturbing the structure of the cell envelope. We note that while depletion of AftB sensitized Mtb to RIF and significantly decreased SL-1 abundance in both cholesterol- and propionate-containing media, the correlation between these two effects was not absolute. AtfB depletion decreased sulfolipid levels to differing degrees under these conditions **(Fig6B)**. This implies that cell wall perturbation might be able to reverse the cholesterol- and lipid-driven RIF resistance phenotype without directly altering lipid abundance.

**Figure 6.**
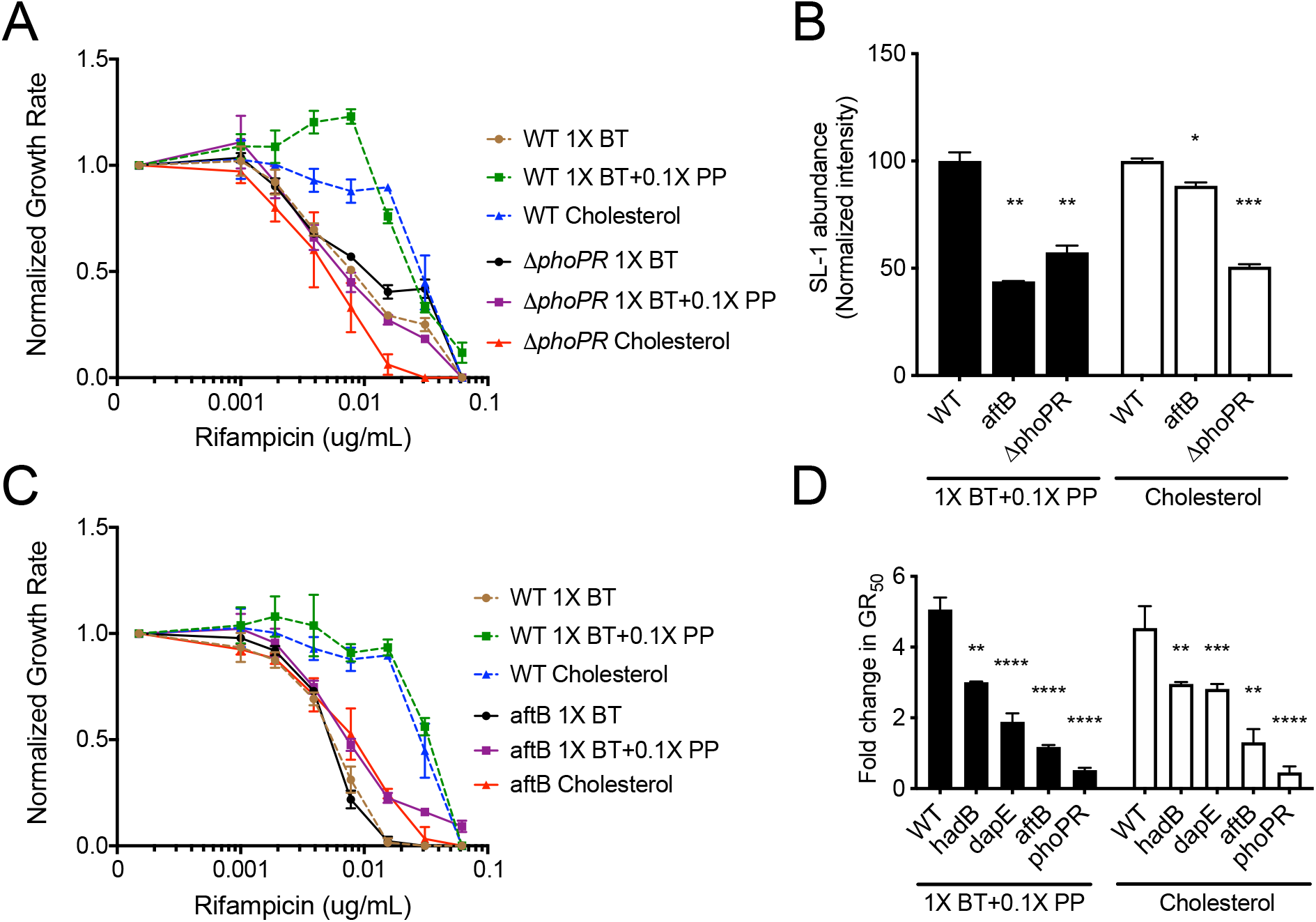
Disruption of the cell envelope reverses propionate and cholesterol catabolism dependent decrease in RIF efficacy. **(A)** Normalized growth inhibition of Δ*phoPR* mutant across increasing concentrations of RIF in minimal media compared to WT. Results shown as means from 3 biological replicates. **(B)** Relative sulfolipid abundances of WT, *aftB* mutant and Δ*phoPR* mutant grown in minimal media with 1X butyrate and 0.1X propionate (1X BT+0.1X PP), and cholesterol. Relative sulfolipid abundances were determined by the combined normalized mass spectra intensities from *m/z* 2000-2800 for each growth condition. MS spectra intensities were normalized using GM2 ganglioside internal standard and cell density. Results shown as means with standard deviations from 2 independent experiments. Significance was calculated using unpaired t-test, \**p*<0.05, \*\**p*<0.01. **(C)** Normalized growth inhibition of *aftB* mutant across increasing concentrations of RIF in minimal media compared to WT. Results shown as means from 3 biological replicates. **(D)** Calculated fold change in GR_50_ values of 1X butyrate and 0.1X propionate (1X BT+0.1X PP) and cholesterol conditions against 1X butyrate alone for WT and mutant strains. Significance was calculated using unpaired t-test, \*\**p*<0.01, \*\*\**p*<0.001, \*\*\*\**p*<0.0001.

### *In vitro* CGI predict drug efficacy *in vivo*

To determine whether CGI identified *in vitro* predict strategies to accelerate bacterial killing in the lung, we infected C57BL/6J mice via the aerosol route with a pooled culture consisting of 3 barcoded WT strains and a number of select hypomorph mutants. We concentrated on genes that either displayed carbon source-independent increase in RIF efficacy (*rpoB, madR*), or mutants that disrupt the peptidoglycan (*murA*) or arabinogalactan (*aftB*) layer of the cells wall and would be expected to reverse the effect of host-derived cholesterol catabolism **(Fig3AB, Fig4BC, S.Fig4)**. To ensure that the animals were treated with a relevant dose of antibiotic, we measured the plasma concentration of RIF in mice and adjusted the dosing to match the 24-hour exposure observed during clinical TB therapy **(Fig7A)** (58). The mice were fed doxycycline chow starting 3 days before infection to repress *sspB* expression in the mutants upon infection, and allow them to grow normally. At 3 weeks post-infection, mice were split to three groups. One group maintained doxycycline for the remainder of the infection without RIF treatment (ND, non-depleted). A second group had doxycycline withdrawn to initiate protein degradation for the remainder of the infection without RIF treatment (D, depleted). The third group had doxycycline withdrawn and two weeks of RIF treatment was administered (D+R, depleted and RIF treated) **(Fig7B)** (59). The antibiotic regimen effectively decreased the total number of lung CFU **(Fig7C)**. Lung homogenates were washed and plated on 7H10 plates with supplementation to recover viable clones. The relative abundance of individual mutants was normalized to the average abundance of three barcoded WT strains for each mouse and fitness changes were calculated by comparing between conditions. Comparing groups D and ND show the relative fitness effects of depletion alone on individual hypomorphs, while examining groups D+R and D show the relative change in fitness due to RIF treatment across depleted samples. Groups ND and D+R were compared to investigate the combined impact of both depletion and RIF treatment.

**Figure 7.**
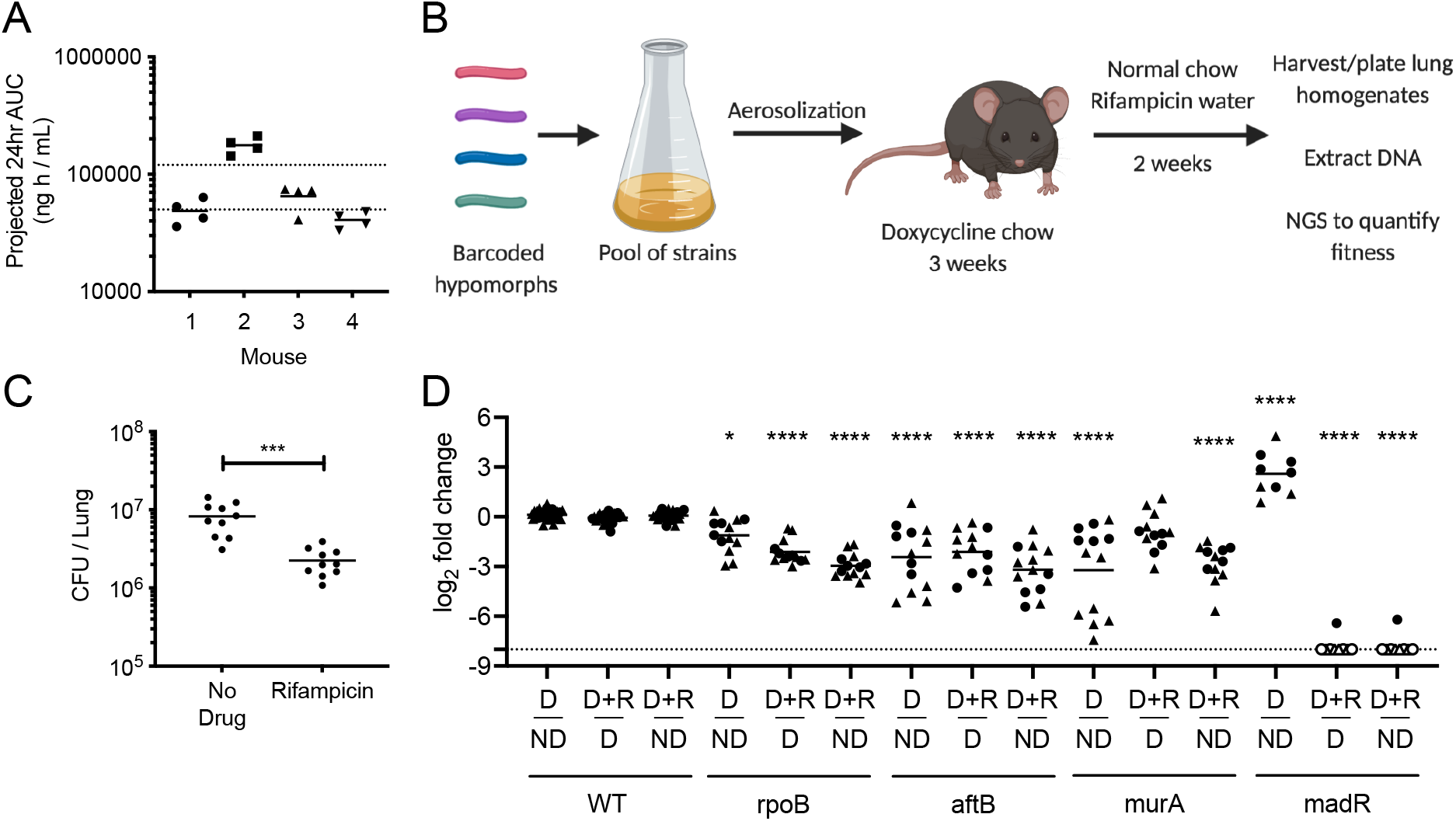
Essential bacterial functions that alter drug efficacy *in vivo*. **(A)** RIF plasma concentrations were measured in mice over 12-hour period 24 hours post RIF administration (0.1g/L). Results shown as Area Under the Concentration (AUC) time profile. Dotted lines indicate RIF plasma range observed during clinical TB therapy (58). **(B)** C57BL/6J mice were infected through the aerosol route with a pooled culture of individual hypomorph mutants and barcoded WT strains. Mice were fed doxycycline chow starting 3 days before infection to 3 weeks post infection. Mice were subsequently switched to normal chow and water with or without RIF (0.1g/L) for 2 additional weeks. Lungs were harvested, homogenized and plated for *Mtb* outgrowth. Upon DNA extraction of grown colonies, chromosomal barcodes were PCR amplified and pooled for Illumina Next Generation Sequencing (NGS). Barcode abundances of individual mutants were normalized to WT strains and analyzed to quantify changes in fitness during RIF treatment. **(C)** RIF treated mice show significant decrease in *Mtb* CFU in lungs compared to untreated controls. Each dot represents a single mouse. Significance was calculated using unpaired t-test, \*\*\**p*<0.001. **(D)** Change in fitness of individual mutants were determined by comparing normalized counts from non-depleted (ND), depleted (D) as well as depleted and RIF treated (D+R) conditions. Depletion of RpoB, AftB and MurA display significant decrease in fitness, shown as Log2 fold change, compared to non-depleted controls. Depletion of RpoB, AftB and MadR show significant decrease in fitness during RIF treatment compared to depleted but untreated controls. Results shown as two independent experiments (circle, triangle) with each dot representing a single mouse. Empty dot indicates that the relative abundance was below the limit of detection (Dotted line). Significance was calculated using Sidak’s multiple comparison test, \**p*<0.05, \*\*\*\**p*<0.0001.

RpoB, AftB and MurA depletion decreased fitness compared to non-depleted controls, while MadR depletion increased relative fitness in the mouse model. Upon RIF treatment, we found that AftB depletion increased bacterial clearance to the same degree as RpoB depletion. Depleting MadR had an even greater effect, and the abundance of this mutant was reduced below our limit of detection in eleven out of twelve animals **(Fig7D)**. The effect of MurA depletion was more variable and its effect on RIF efficacy was not statistically significant. In sum, the reproducible effects of RpoB, AtfB, and MadR on RIF efficacy show that CGI identified *in vitro* can accelerate bacterial clearance during infection, and highlight the importance of the environment for defining relevant interactions.

## Discussion

Optimizing combination therapy is critical to improving TB treatment. While efficient strategies for predicting and quantifying drug-drug interactions under *in vitro* conditions have been developed, the impact of bacterial environment on these interactions has not been determined (27, 28). As a result, it remains unclear how well these *in vitro* data predict the effects of combination therapy in the infection environment. In this study, we modeled drug-drug interactions using a newly developed genetic hypomorph library to assess the impact of bacterial carbon metabolism on CGI. This work revealed that CGI with distinct drugs are differentially sensitive to the environment, suggesting tailored approaches to optimization.

While global analysis of our chemical-genetic-environmental dataset suggested that drug mechanism and environment both play critical roles in shaping chemical-genetic interactions, all drugs were not equally affected by condition. In particular, the individual effect of the fluoroquinolone, MOX, was unaffected by carbon source, and this relative indifference to the environment was also observed for CGI with MOX. These observations contrast with RIF and INH, where carbon source plays an important role in determining the MIC of the drug alone, as well as shaping chemical-genetic interaction profiles. While we can only speculate on the mechanistic basis for these differences between drugs, we found that the cholesterol-dependent decrease in RIF sensitivity is due to the effect of carbon source on cell envelope structure that is likely to alter RIF penetration. Similarly, the activation of the prodrug, INH, is affected by the redox state of the cell (60), another process that is impacted by carbon catabolism. In contrast, MOX is not a prodrug and we speculate that it may not be limited by cellular penetration as strongly as RIF. More practically, these observations suggest that drug-drug interactions with MOX may be particularly robust to changes in condition, and therefore may translate well between *in vitro* and *in vivo* conditions.

Complex drug-drug interactions may both increase or decrease sensitivity. Our dataset found potential antagonistic interactions that may arise with different TB antibiotics. For example, the inhibition of Ndh, a NADH-dehydrogenase, has previously been shown to decrease INH efficacy and was also identified in our dataset (37). Antagonistic interactions have important implications for TB therapeutics due to the reliance on multi-drug regimens, and our study provides a global assessment of these effects.

While RIF interactions were complex and condition-dependent compared to MOX, we found that the cholesterol-specific effects translate well to the infection environment and may be particularly relevant to treatment. As reduced RIF exposure limits the efficacy of this critical component of our standard regimen, even small increases in MIC during infection could have a significant effect on bacterial killing. A number of observations indicate that RIF efficacy is limited by similar mechanisms *in vivo* and in cholesterol-containing media. Firstly, the increase in branched-chain lipid abundance that is associated with reducing RIF efficacy in cholesterol- or propionate-containing media has also been observed during infection, where similar carbon sources are utilized (16, 53). Furthermore, genetic perturbation of the same cell wall synthetic enzymes sensitized Mtb to RIF in both conditions. This effect may underlie the increased cell-associated RIF upon simultaneous INH and ethambutol treatment *in vitro* (61–63). While the importance of cell wall architecture and cell envelope lipid production suggests that cellular permeability to RIF may be altered during growth on cholesterol, we note that lipid synthesis also has important effects on central carbon metabolic pathways that could influence antibiotic susceptibility (15). Furthermore, the mixture of carbon sources available in different host microenvironments remains ill-defined, and cholesterol may not be the only macronutrient capable of limiting RIF efficacy. Additional enzymes identified in this study may also be attractive targets for further drug development, such as AftB, a arabinofuranosyltransferase distinct from EmbA and EmbB that are targets of ethambutol (47). Similarly, enzymes in the DAP (diaminopimelic acid) biosynthesis pathway produce precursors for the synthesis of peptidoglycan and lysine (64), and the loss of either sensitizes the cell to RIF.

Using a chemical-genetic system, we determined that cellular environment plays an important role in shaping CGIs, and that even the use of relatively simple *in vitro* culture conditions can identify interactions that are relevant during infection. These studies provide a tractable system that can incorporate more complex culture systems, or even cellular models, to identify additional *in vivo*-relevant interactions. Expanding this CGI atlas promises to elucidate the processes limiting drug efficacy during infection and guide drug development and regiment optimization efforts.

## Material and methods

### Bacterial cultures

*M. tuberculosis* (H37Rv) cells were cultured in Middlebrook 7H9 medium supplemented with 10% oleic acid-albumin-dextrose-catalase (OADC), 0.5% glycerol, and 0.05% Tween-80, or in minimal medium supplemented with 0.1% Tyloxapol and variable levels of glycerol, acetate, butyrate, propionate, or cholesterol up to 0.1%. Minimal medium was made as previously described but with ferric chloride (100uM) replacing ferric ammonium citrate (51).

Cells were also cultured on 7H10 agar medium supplemented with 10% OADC and 0.5% glycerol. Streptomycin (20ug/mL) was supplemented when necessary. Anhydrotetracycline (ATC, 500ng/mL) was supplemented periodically until cultures reach exponential growth. ATC was removed from cultures prior to antibiotic growth inhibition assays by washing in PBS supplemented with 0.1% Tyloxapol. Δ*phoPR* mutant was generated as described previously (65).

### Multiplexed library screening

Library pools of hypomorph mutants were prepared as described previously (32). Pools were initially cultured in Middlebrook 7H9 medium supplemented with 10% oleic acid-albumin-dextrose-catalase (OADC), 0.5% glycerol, and 0.05% Tween-80. Cultures were supplemented with Anhydrotetracycline (ATC, 500ng/mL) periodically until cultures reach exponential growth. ATC was removed from cultures by washing in PBS supplemented with 0.1% Tyloxapol prior to inoculation in 96-well plates to allow induction of SspB. Pools were grown in minimal medium with 0.1% glycerol, 0.1% acetate, or 0.1% cholesterol on 96-well plates for 2 weeks. Isoniazid and moxifloxacin were used at 1ug/mL and rifampicin was used at 0.062 ug/mL. Antibiotics were added to select wells and serially diluted in a 2-fold manner prior to inoculation. An untreated growth condition was included for each study. Samples were inoculated at OD600 0.01 and growth was monitored using OD600. Upon completion, 96-well plates were heat-inactivated at 85 degrees for 2 hours. Barcodes were PCR amplified as described previously (32). Individual libraries were mixed with 1:1 20% DMSO and heated for 10 minutes at 98 degrees prior to PCR reaction. Amplified barcodes were purified using SPRI-based purification methods and sequenced using Next-generation sequencing methods. Sequence alignment and analysis were conducted using Bowtie software package with index mismatch set to 2 bases and barcode mismatch set to 1 base. Relative abundance of every mutant was calculated as mean ratio of barcode abundance of mutant relative to total barcode abundance of library. Log2FC of a mutant was calculated as the log2 of the ratio of the mean relative abundances in a given antibiotic condition relative to its untreated control. Hierarchical clustering and principal component analysis were conducted using ClustVis (66). Hierarchical clustering was applied to vectors of Log2FC of each gene across all conditions. PCA was performed on relative abundances across all conditions. Statistical significance was determined by unpaired *t*-test with Benjamini-Hochberg multiple testing correction.

### Antibiotic growth inhibition assay

*Mtb* cells were grown in minimal medium with 0.1% glycerol, 0.01% acetate, 0.01% butyrate, 0.01% propionate, or 0.01% cholesterol on 96-well plates. Isoniazid and moxifloxacin were used at 1ug/ml and rifampicin was used at 0.062 ug/mL. Antibiotics were added to select wells and serially diluted in a 2-fold manner prior to inoculation. Untreated condition was included for each study. Cells were inoculated at OD600 0.01 and growth was monitored using OD600 until exponential phase (OD600 0.20-0.25). Antibiotic efficacy was determined using growth rate inhibition as done previously (24). The exponential growth constant (*k*) value was calculated by plotting an exponential curve over all growth points. The *k*-value of each antibiotic concentration was normalized to the *k*-value of the no-drug control. The GR_50_ value was defined as the concentration of antibiotic that resulted in 50% decrease in growth rate. L-lysine (Sigma) was added to growth conditions to final concentration of 3mM.

### Lipid extraction and mass spectrometry

*Mtb* cells were grown in minimal medium with 0.01% butyrate, propionate or cholesterol. Cells were inoculated at OD_600_ 0.1 and grown to final OD_600_ of 0.7. Cells were pelleted and heat-inactivated at 85 degrees for 45 minutes. Cells were washed in 10% glycerol to remove residual detergents from growth media. Lipid extraction was conducted with methyl tert-butyl ether (MTBE) as described previously (67). 1.5mL methanol and 5mL MTBE were mixed with the cell pellet and placed into a glass tube with a Teflon-lined cap. The mixture was incubated at room temperature for 5 hours while shaking. Phase-separation was achieved by adding 0.75mL water to the tubes, incubating for 10 minutes at room temperature, then centrifuging at 1000g for 10 minutes. The 2.5mL of the top phase was collected and transferred to a fresh glass tube with Teflon-lined cap. Samples were dried using a nitrogen evaporator and stored in −20 degrees prior to injection to mass spectrometer. After evaporation of MTBE, 300 μL of 2:1 methanol:chloroform was added to each tube followed by vigorous vortex. Solubilized lipids were then transferred to a new, pre-weighed tube, dried using a nitrogen evaporator, and then tubes were re-weighed to determine the lipid mass in each sample. Then 1.5 μL of 100 ng/μL N-omega-CD3-octadecanoyl monosialoganglioside GM_2_ (Matreya, State College, PA) was added to each sample for normalization and samples were reconstituted in 300 μL of 2:1 methanol: chloroform for mass spectrometry analysis.

Samples were analyzed by direct infusion on a syringe pump to an Orbitrap Velos Pro (Thermo Fisher Scientific, Waltham, MA) mass spectrometer operating in the negative electrospray ionization mode. Mass spectra were acquired at a flow rate of 10 μL/min for 2 minutes in two different mass ranges, *m/z* 200-2000 and *m/z* 300-3000, acquiring two replicates for each range using a resolution of 30,000 (*m/z* 200), an AGC target population of 5e^5^ and a maximum ion injection time of 100 ms. Data were analyzed in Xcalibur 2.2 (Thermo Scientific). Briefly, mass spectra were averaged over the entire acquisition range (0-2 minutes), producing one averaged spectrum for each analysis. Peak lists were exported in Excel and intensity data was normalized to the intensity of the GM2 ganglioside spike.

### Plasma pharmacokinetic analysis

C57BL/6J mice were purchased from Jackson Laboratories. Housing and experimentation were done in accordance with the guidelines set by the Department of Animal Medicine of University of Massachusetts Medical School and Institutional Animal Care and Use Committee and adhered to the laws of the United States and regulations of the Department of Agriculture. Eight-to 12-week mice were administered 0.1g/L RIF through drinking water for three days. Blood was collected during a 12-hour period 24 hours post RIF administration, kept on ice, centrifuged at approximately 2500 x g for 5 minutes. After centrifugation, plasma was collected and stored at −80°C until analysis.

Neat 1 mg/mL DMSO stocks for RIF were serial diluted in 50/50 acetonitrile/water to create neat spiking stocks. Standards and quality control (QC) samples were created by adding 10 μL of spiking stock to 90 μL of drug free plasma. Ten microliters of control, standard, QC, or study sample were added to 100 μL of acetonitrile/methanol 50/50 protein precipitation solvent containing 20 ng/mL RIF-d8. Extracts were vortexed for 5 minutes and centrifuged at 4000 RPM for 5 minutes. 100 μL of supernatant was combined with 5 μL of 75 mg/mL ascorbic acid to stabilize RIF. 100 μL of mixture was combined with 100 μL of Milli-Q water prior to HPLC-MS/MS analysis. Mouse control plasma (K2EDTA) was sourced from Bioreclamation. Mouse control lungs were collected in house. RIF was sourced from Sigma Aldrich and RIF-d8 was purchased from Toronto Research Chemicals.

LC-MS/MS analysis was performed on a Sciex Applied Biosystems Qtrap 6500+ triple-quadrupole mass spectrometer coupled to a Shimadzu Nexera X2 UHPLC system to quantify each drug in plasma. Chromatography was performed on a Agilent SB-C8 (2.1×30 mm; particle size, 3.5 μm) using a reverse phase gradient. Milli-Q deionized water with 0.1% formic acid was used for the aqueous mobile phase and 0.1% formic acid in acetonitrile for the organic mobile phase. Multiple-reaction monitoring of precursor/product transitions in electrospray positive-ionization mode was used to quantify the analytes. Sample analysis was accepted if the concentrations of the quality control samples were within 20% of the nominal concentration. The compounds were ionized using ESI positive mode ionization and monitored using masses RIF (823.50/791.60) and RIF-d8 (831.50/799.60). Data processing was performed using Analyst software (version 1.6.2; Applied Biosystems Sciex).

### *In vivo* antibiotic susceptibility

C57BL/6J mice were purchased from Jackson Laboratories. Housing and experimentation were done in accordance with the guidelines set by the Department of Animal Medicine of University of Massachusetts Medical School and Institutional Animal Care and Use Committee and adhered to the laws of the United States and regulations of the Department of Agriculture. Eight-to 12-week mice were infected with pools of strains at equal ratios through the aerosol route (500 to 1000 CFU/mouse). Mice were fed doxycycline-containing chow (Purina 5001 with 2000 ppm doxycycline, Research Diets C11300-2000i) starting 3 days pre-infection to 21 days post-infection. At 21 days post-infection, 0.1g/L rifampicin was administered through drinking water for 14 days. At 35 days post-infection, mice were sacrificed, spleen and lungs were isolated and homogenized, and CFU was determined by plating dilutions on 7H10 agar with 50ug/mL streptomycin and 500ng/mL ATC. For library recovery, approximately 1 million CFU per mouse were plated on 7H10 agar with 50ug/mL streptomycin and 500ng/ml ATC.

Genomic DNA was extracted and normalized as done previously (35) and sequenced using multiplex PCR methods. Sequence alignment and analysis were conducted using Bowtie software package as described above. Relative abundance of every mutant was calculated as mean ratio of barcode abundance of mutant relative to the average barcode abundance of three barcoded H37Rv strains. Log2FC of a mutant was calculated as the log2 of the ratio of the mean relative abundances of rifampicin treated mice relative to the untreated mice. Statistical significance was determined by unpaired *t*-test.

## Acknowledgements

We are thankful to the members of the Sassetti lab for both technical assistance and helpful discussion. We thank Sovie Lavalette-Levy and Curtis Engelhart for technical help. This work was supported by the Office of the Assistant Secretary of Defense for Health Affairs through the Peer Reviewed Medical Research Program, Focused Program Award, under award no. W81XWH-17-1-0692. Opinions, interpretations, conclusions, and recommendations are those of the author and are not necessarily endorsed by the Department of Defense. The work was additionally supported by the NIH (AI095208). Model figures were created with BioRender.com

