## Supplemental Figures for "Chemical-genetic interaction mapping links carbon metabolism and cell wall structure to tuberculosis drug efficacy"

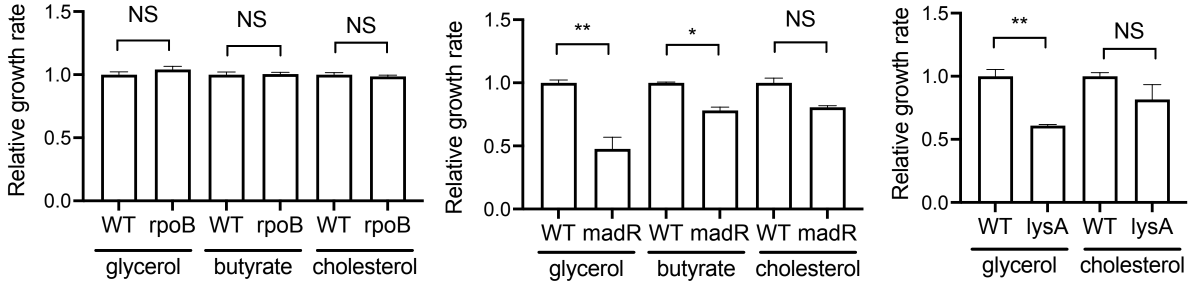


**Supplementary Figure 1. Relative growth rates of hypomorph mutants that show condition-independent interactions with RIF.** Relative growth rates of WT and *rpoB* **(left)**, *madR* **(center)**, *lysA* **(right)** hypomorph mutant during untreated growth in respective carbon sources. Growth rate of hypomorph mutant was normalized to the WT growth rate from the respective carbon source. Results shown as means from 3 biological replicates with standard deviations. Significance was calculated using unpaired t-test, *$p$<0.05, **$p$<0.01, Not Significant (NS).

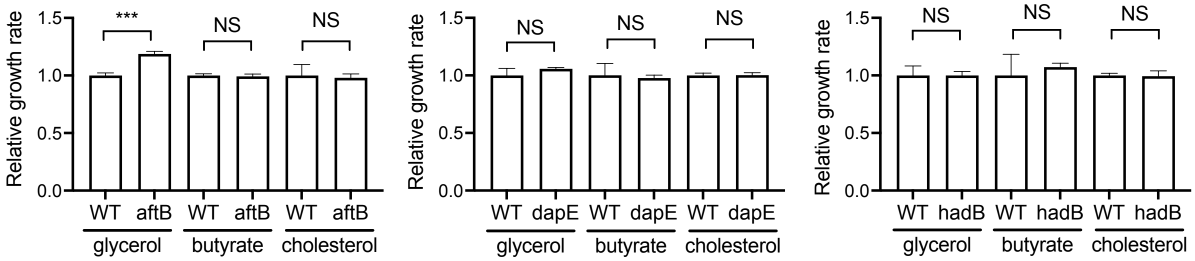


**Supplementary Figure 2. Relative growth rates of hypomorph mutants that show condition-dependent interactions with RIF.** Relative growth rates of WT and *aftB* **(left)**, *dapE* **(center)**, *hadB* **(right)** hypomorph mutant during untreated growth in respective carbon sources. Growth rate of hypomorph mutant was normalized to the WT growth rate from the respective carbon source. Results shown as means from 3 biological replicates with standard deviations. Significance was calculated using unpaired t-test, ***$p$<0.001, Not Significant (NS).

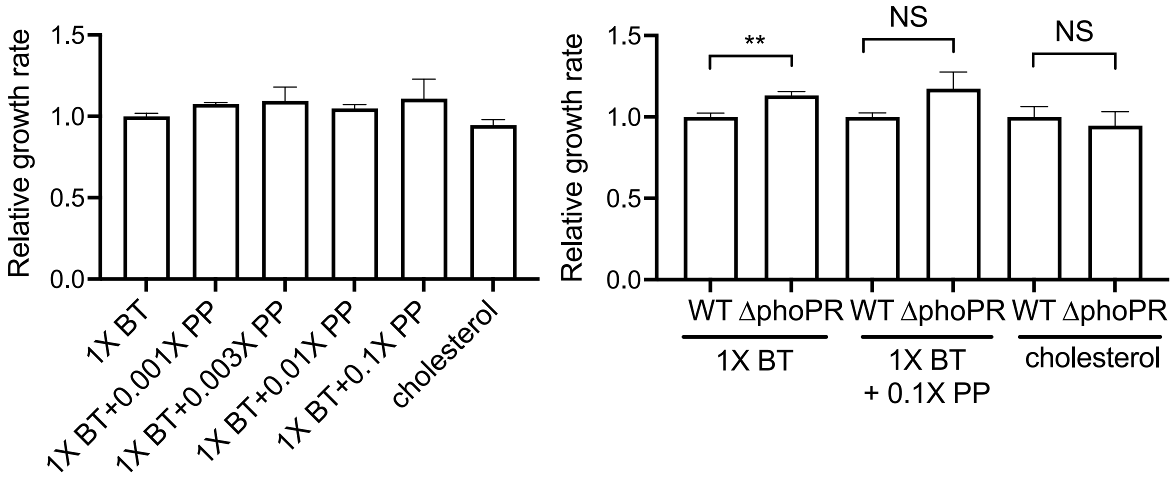


**Supplementary Figure 3. Relative growth rates of WT and Δ*phoPR* mutant from different carbon sources. (left)** Relative growth rates of WT during untreated growth in different carbon sources. Growth rates of propionate supplemented conditions (PP) and cholesterol were normalized to the butyrate (BT) growth rate. No significance was seen between different conditions. Results shown as means from 3 biological replicates with standard deviations. Significance was calculated using unpaired t-test. **(right)** Relative growth rates of WT and Δ*phoPR* mutant during untreated growth in respective carbon sources. Growth rate of Δ*phoPR* mutant was normalized to the WT growth rate from the respective carbon source. Results shown as means from 3 biological replicates with standard deviations. Significance was calculated using unpaired t-test, ***p*<0.01, Not Significant (NS).

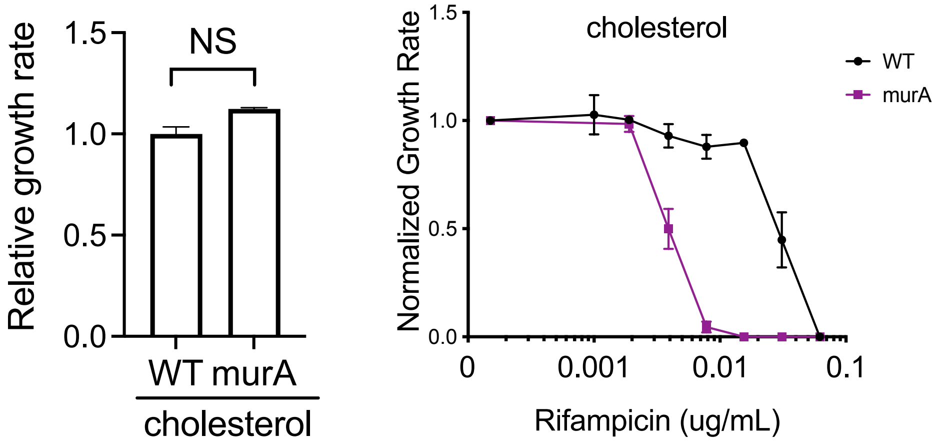


**Supplementary Figure 4. MurA hypomorph mutant shows altered RIF efficacy during cholesterol growth conditions. (left)** Relative growth rates of WT and murA hypomorph mutant during untreated growth in respective carbon sources. Growth rate of hypomorph mutant was normalized to the WT growth rate from the respective carbon source. Results shown as means from 3 biological replicates with standard deviations. Significance was calculated using unpaired t-test, Not Significant (NS). **(right)** Normalized growth inhibition of WT and MurA hypomorph mutant across increasing concentrations of RIF in minimal media with cholesterol as the sole carbon source. Depletion of MurA (purple) show decreased RIF MIC compared to WT (black) during cholesterol growth conditions. Results shown as means from 3 biological replicates with standard deviations.
